# The Anadromous Hickory Shad (Clupeiformes: Clupeidae, *Alosa mediocris* [Mitchill 1814]): Morphometric and Meristic Variation

**DOI:** 10.1101/716183

**Authors:** Jordan P. Smith, Michael S. Brewer, Roger A. Rulifson

## Abstract

The anadromous Hickory Shad *Alosa mediocris* (Mitchill, 1814) (Clupeiformes: Clupeidae) is reviewed, specifically regarding morphometric and meristic variation. Despite its long history as recognized species, few descriptions of Hickory Shad morphometric and meristic characters exist in the literature. Most authors of the historic literature have failed to provide capture location for specimens, analyze large numbers of Hickory Shad, or document how morphometric and meristic characters of the species vary spatially. To address this information gap, a total of 717 mature Hickory Shad were collected from 23 different locations in Maryland, Delaware, Virginia, North Carolina, South Carolina, Georgia, and Florida using electroshocking, gill net, or rod and reel. All specimens were frozen, thawed, and 17 morphometric characters and four meristic characters were examined; a random subset (n = 463) were analyzed for an additional four meristic counts of gill rakers. Overall specimens ranged from 206-389 mm SL with a mean + SD of 278.41 + 27.69 mm, 232-435 mm FL with a mean of 310.98 + 30.35 mm, and 272-508 mm TL with a mean of 365.62 + 35.52 mm. The linear relationships between FL and TL, and FL and SL, were investigated and found to be: TL = 1.169*FL + 1.660 (n=705, r^2^=0.995) and SL = 0.909*FL - 4.274 (n=717, r^2^=0.992). Substantial differences in character means for many morphometric measurements were found between male and female specimens, suggesting strong sexual dimorphisms relating to shape. However, meristic characters did not show differences in character means by sex. No one morphometric measurement could distinguish Hickory Shad from other morphologically similar clupeids, but the meristic count of gill rakers on the lower limb of the first arch were important to separate Hickory Shad (19-22) from American Shad *A. sapidissima* (Wilson, 1811), Alewife *A. pseudoharengus* (Wilson, 1811), and Blueback Herring *A. aestivalis* (Mitchill, 1814).

## Introduction

No published study has examined and described an extensive set of morphometric and meristic characters of the Hickory Shad *Alosa mediocris* (Mitchill, 1814) (Clupeiformes: Clupeidae). The initial description of Hickory Shad lacked some critical information, indicating that this was a species unknown to the system and proceeded to describe it from “fresh specimens.” Unfortunately there is no reference to the capture location of the fish nor quantity examined. It is possible that the description could have been based from one or several individuals. We speculate that the likely watershed from which Mitchill collected his specimen(s) was the Hudson River due to its close proximity to Columbia University.

Few records exist of Mitchill’s early attempts to describe New York fauna, including *A. mediocris*. Perhaps Professor Mitchill took students to the shores of the Hudson River to observe fauna from pulling small seines; unless more early writings of Professor Mitchill are discovered, the locations and manner of these ichthyological collections will remain unknown. One or more of those specimens collected was an undescribed species of “Shad”, which he presumably took back to his laboratory for examination and decided the specimen(s) fit within the family Clupeidae. Mitchill proceeded to designate the species *Clupea mediocris –* the “Staten Island Herring”. In a presumably similar manner, Mitchill also described 11 other new species during that era (including what is now known as *Alosa aestivalis* (Mitchill, 1814), the Blueback Herring) although all 12 new “Mitchillian” species, including the current-day Hickory Shad and Blueback Herring, were placed in different genera by subsequent authorities [1].

Unfortunately, the original description of the Hickory Shad contained only a sparse description of the anatomical features. Mitchill [2] included basic descriptions of the fish shape, color, size, and meristic counts for branchiostegal, pectoral, ventral, anal, dorsal, and caudal fin rays, but he did not include any information on morphological measurements or ratios of size between various body features. Interestingly, many researchers describing the few characteristics of this species did so citing other investigators, who in turn cited Mitchill [2]. Therefore, little additional meristic or morphological information has been recorded for the species since the original description, over 200 years ago.

In addition, no record can be located of the holotype, nor where or when the specimen was collected. During this time of budding taxonomy in America, it was neither common nor required to keep holotype specimens for newly described species. Other taxonomists after Mitchill revised the taxonomic status of the Hickory Shad. Notably, the genus *Alosa* was divided into three genera by Regan [3] in 1917: *Alosa*, *Caspialosa*, and *Pomolobus*; the Hickory Shad was classified under the genus *Pomolobus* along with the Alewife and the Blueback Herring [4]. Later work by Bailey [5] and Svetoviodov [6] led to synonymizing the genera *Pomolobus* and *Caspialosa* with the genus *Alosa*, thereby changing the scientific name of Hickory Shad from *Pomolobus mediocris* to *A. mediocris* (Mitchill, 1814) [4].

Mansueti [7] examined the hypothesis that the Hickory Shad might be a hybrid between the American Shad *Alosa sapidissima* (Wilson, 1811) and one of the River Herrings, the Alewife *Alosa pseudoharengus* (Wilson, 1811) or the Blueback Herring *A. aestivalis.* He concluded that hybridization was unlikely and “not substantiated by any reliable evidence” [7]. Around this time, a few fish culturists experimented in hatcheries and actively pursued creating hybrids involving Hickory Shad and River Herring, though none of these attempts were successful [7].

The objective of this manuscript is to fully describe the various anatomical features, including meristic counts and morphological measurements, of the Hickory Shad across its range. The Hickory Shad is considered an understudied fish species though it spawns in rivers on the United States Eastern Seaboard from the Schuylkill River in the Delaware River watershed [8] to the St. Johns River in Florida [9]. The northern range limit of Hickory Shad spawning populations is not precisely known; early authors hypothesized spawning as far north as Maine [10]. A spawning population is suspected in Wethersfield Cove in the Connecticut River near Wethersfield, Connecticut, but evidence is lacking; adult Hickory Shad have been collected from that area during spring (Ken Sprankle, USFWS, personal communication). Rulifson [11] reported that Connecticut is the northernmost state having a presence of Hickory Shad based on responses to questionnaires by respective state fisheries biologists. It is possible some of these northern accounts of Hickory Shad are either misidentifications with morphologically similar species, such as the American Shad *A. sapidissima,* or possibly wandering Hickory Shad collected in bays or the Atlantic Ocean, but not actively spawning. The Hickory Shad is a schooling species of the family Clupeidae and utilizes the life history strategy of anadromy, entering coastal freshwater between February and June to spawn; the higher latitudes correspond to later dates of entry into freshwater [12].

Relatively few authors have included morphometric and meristic values for Hickory Shad [13], [14], [15], [10], [16], [17], [18] but none investigated how these characters vary spatially. Most previous studies fail to provide capture location(s) for the specimens examined and cover many fewer characters than the present study. Furthermore, some authors provide only one value for various meristic counts and morphometric measurements, when in reality there is often considerable variation. No published study has described Hickory Shad specimens across such a large latitudinal gradient, covering the majority of the species range. Similar studies have been undertaken for the American Shad [19], Alewife, and Blueback Herring [20].

Historically, morphometric and meristic analyses of fish have been valuable tools for early ichthyologists and naturalists alike [21]. Starting in 1894, the Royal Society of the United Kingdom created the “Committee for Conducting Statistical Inquiries into the Measurable Characters of Plants and Animals.” One of the committee’s chief tasks was to investigate morphometric variation in Atlantic Herring *Clupea harengus* (Linneaus, 1758) [22]. Analysis of morphological and meristic characters of fish is straightforward, cost-efficient, and an often-used tool to identify and differentiate fish species, stocks, and populations [23].

## Methods

Hickory Shad specimens were collected during the 2016 and 2017 spawning runs from the Susquehanna and Patapsco rivers, Maryland; the Nanticoke River, Delaware; the Rappahannock, Appomattox, and James rivers, Virginia; the Chowan River headwaters (Meherrin, Nottaway, and Blackwater), also in Virginia; the Roanoke, Cashie, Pungo, Pamlico, Tar, Neuse, New, and Cape Fear rivers, North Carolina; Pamlico Sound, also in North Carolina; the Waccamaw and Santee rivers, South Carolina; the Altamaha River, Georgia, and the St. Johns River, Florida (Table 1). In addition, a few specimens (n=5) were obtained from the Atlantic Ocean close to shore, near Wrightsville Beach, North Carolina. Relative location of rivers as well as collection sites are depicted in Figure 1. All specimens were collected from the different locations by recreational angling (i.e., rod and reel), gill net, or electrofishing. Specimens from rivers outside of North Carolina were collected and donated to this study by the respective state or federal fisheries agencies. North Carolina fish came from the North Carolina Wildlife Resources Commission (NCWRC) or the North Carolina Division of Marine Fisheries (NCDMF). Additional sampling was conducted by the Rulifson Lab with electrofishing and rod and reel (NC Scientific Collection Permit Number 17-SFC00133; East Carolina University AUP #D330).

**Table 1.** List of states and river (north to south), sex, and total number of Hickory Shad collected in 2016 and 2017.

| State | River | Sex |  | Unknown | Total |
| --- | --- | --- | --- | --- | --- |
|  |  | Female | Male |  |  |
| Maryland | Susquehanna R. | 13 | 9 |  | 22 |
| Maryland | Patapsco R. | 11 | 39 |  | 50 |
| Delaware | Nanticoke R. | 16 | 6 |  | 22 |
| Virginia | Rappahannock R. | 23 | 21 | 3 | 47 |
| Virginia | Appomattox R. | 25 | 25 |  | 50 |
| Virginia | James R. | 26 | 37 | 2 | 65 |
| Virginia | Chowan R. (Meherrin) |  | 1 |  | 1 |
| Virginia | Chowan R. (Nottaway) | 7 | 11 |  | 18 |
| Virginia | Chowan R. (Blackwater) | 13 | 11 | 1 | 25 |
| North Carolina | Roanoke R. | 21 | 23 |  | 44 |
| North Carolina | Cashie R. | 17 | 17 |  | 34 |
| North Carolina | Pamlico Sound | 63 | 29 | 2 | 94 |
| North Carolina | Pungo R. |  | 2 | 1 | 3 |
| North Carolina | Pamlico R. | 39 | 24 | 1 | 64 |
| North Carolina | Tar R. | 31 | 20 | 1 | 52 |
| North Carolina | Neuse R. | 14 | 30 | 3 | 47 |
| North Carolina | New R. | 2 | 2 | 6 | 10 |
| North Carolina | Atlantic Ocean* | 3 |  | 2 | 5 |
| North Carolina | Cape Fear R. | 5 | 13 |  | 18 |
| South Carolina | Waccamaw R. | 7 |  |  | 7 |
| South Carolina | Santee R. | 2 | 4 |  | 6 |
| Georgia | Altamaha R. | 26 | 4 |  | 30 |
| Florida | St. Johns R. (Wekiva) | 1 | 2 |  | 3 |
|  |  | **365** | **330** | **22** | **717** |

\*denotes non-river or sound sampling location

**Figure 1.**
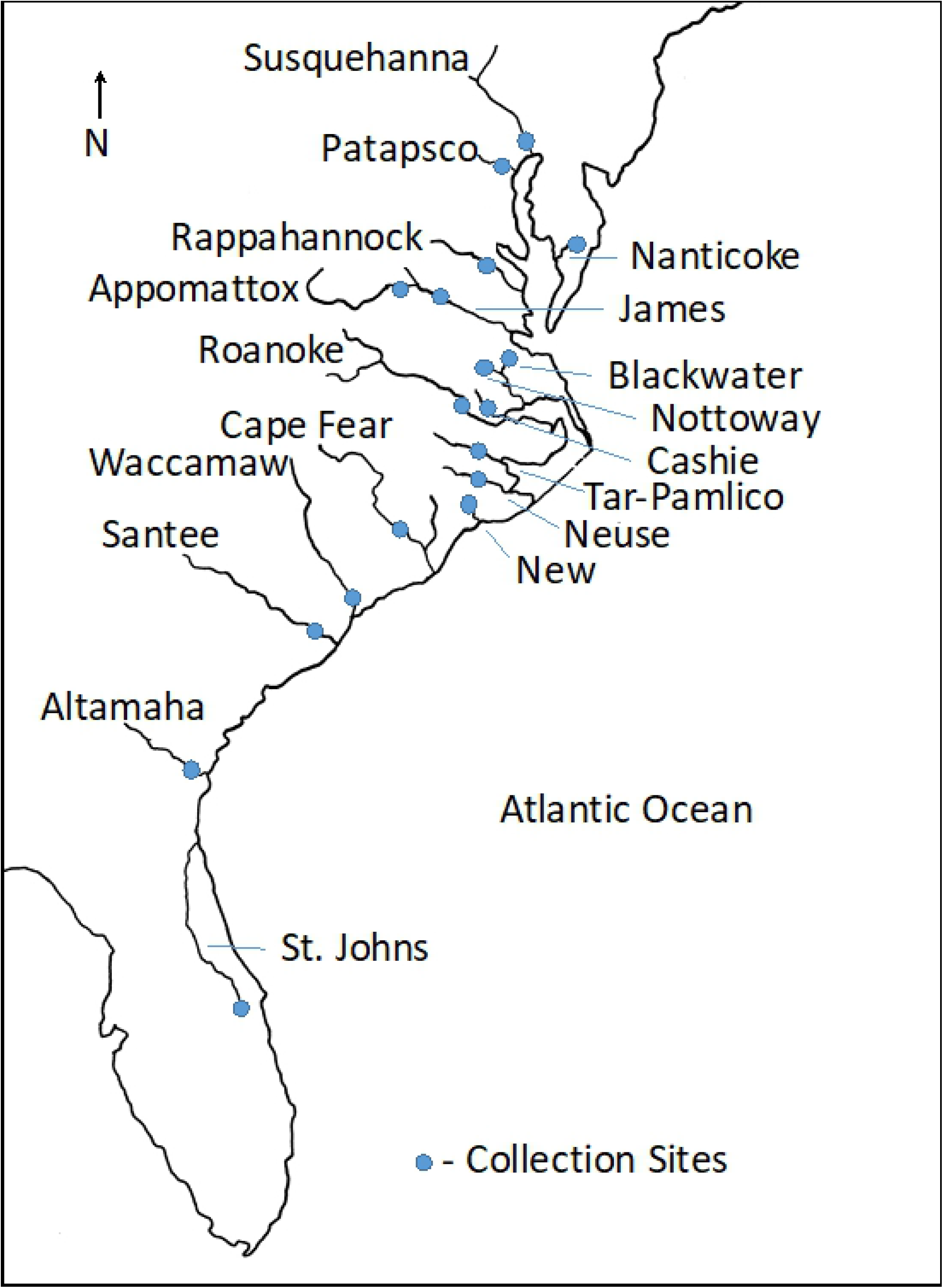
Map showing relative location of rivers included in this study as well as collection sites of Hickory Shad. Revised after Melvin et al. [28].

Initially all specimens were frozen in water to minimize freezer burn and fin breakage, and then eventually transferred to the Rulifson Lab at East Carolina University (ECU) for examination. Once received or collected, fish were identified to species based on projection of the lower jaw beyond the maxilla (as opposed to the American Shad, for which the lower jaw inserts into a slot in the maxilla), weighed to the nearest 0.01 g, bagged individually without water, and given a unique identification number. After this step the fish were placed in freezers (−20°C or −0°C) on the ECU campus until analysis. Specimens were removed from the freezer and slowly allowed to thaw. A small tissue sample was taken from the dorsal fin, which was then placed in 95% ethanol (ETOH) and stored in a −80°C freezer for later genetic analysis.

A total of 17 morphometric measurements and 4 meristic characters (Table 2) were recorded generally following the methods outlined by Hubbs and Lagler [24]. All measurements were straight line distances from point to point on the left side of the body unless there was physical damage: standard length (SL) -- distance between most anterior portion of the head (lower jaw) to the last vertebrae; fork length (FL) -- the distance between the lower jaw to the fork of the caudal tail; total length (TL) -- the greatest distance between lower jaw and end of caudal fin when the caudal rays are pinched together; lower lip to nose (LLN) -- the distance of the projecting lower jaw to maxilla; snout to anal length (SAL) -- the distance between lower jaw and the anus; body depth (BD) -- greatest depth distance between anterior to dorsal fin and anterior of the ventral fin; head length (HL) -- the distance from lower jaw to the most distant point of the operculum (including membrane); eye length (EL) -- the greatest distance of the orbit; snout length (SNL) -- the distance from the most anterior point of the upper lip to the anterior margin of the orbit; head width (HW) -- the distance (width) across the head where the preopercle ends; interorbital width (IOW) -- distance between the eyes at the top of the cranium; maxillary length (ML) -- the distance from the tip of the upper jaw to the distal end of the maxillary; fin length dorsal base (FLD) -- the greatest distance of the structural base between the origin and insertion of the dorsal fin when the fin is erect; fin length anal base (FLA) -- the greatest distance of the structural base between the origin and insertion of the anal fin when the fin is erect; longest ray dorsal fin (LRD) -- the distance from the structural base of the dorsal fin to the tip of the longest ray; longest ray pectoral fin (LRP) -- distance from the structural base of the pectoral fin to the tip of the longest ray; longest ray ventral (pelvic) fin (LRV) -- distance from the structural base of the ventral fin to the tip of the longest ray; longest ray anal fin (LRA) -- distance from the structural base of the anal fin to the tip of the longest ray when the fin is erect. A Hickory Shad illustration (Figure 2) depicts how most morphometric measurements were taken. IOW and HW were omitted on the illustration since they are width measurements and cannot be accurately depicted. The standard length, total length, and snout-to-anal length were measured to the nearest mm; all other measurements were taken by using Fisherbrand “Traceable” digital calipers (model number 06-644-16) to the nearest 0.01 mm.

**Table 2.** Morphometric measurements and meristic counts analyzed, and acronyms used in this study.

| **Morphometric** | **Acronym** | **Meristic** | **Acronym** |
| --- | --- | --- | --- |
| Standard Length | SL | Posterior Ventral Scutes | PVS |
| Fork Length | FL | Anterior Ventral Scutes | AVS |
| Total Length | TL | Scale Rows | SR |
| Lower Lip-Nose | LLN | Longitudinal Scale Rows | LSR |
| Snout-to-Anal Length | SAL | Left Gill Raker Upper | L-GRU |
| Body Depth | BD | Right Gill Raker Upper | R-GRU |
| Head Length | HL | Left Gill Raker Lower | L-GRL |
| Eye Length | EL | Right Gill Raker Lower | R-GRL |
| Snout Length | SNL |  |  |
| Head Width | HW |  |  |
| Interorbital Width | IOW |  |  |
| Maxillary Length | ML |  |  |
| Fin Length-Dorsal Base | FLD |  |  |
| Fin Length-Anal Base | FLA |  |  |
| Longest Ray Dorsal Fin | LRD |  |  |
| Longest Ray Left Pectoral Fin | LRP |  |  |
| Longest Ray Left Ventral Fin | LRV |  |  |
| Longest Ray Anal Fin | LRA |  |  |

**Figure 2.**
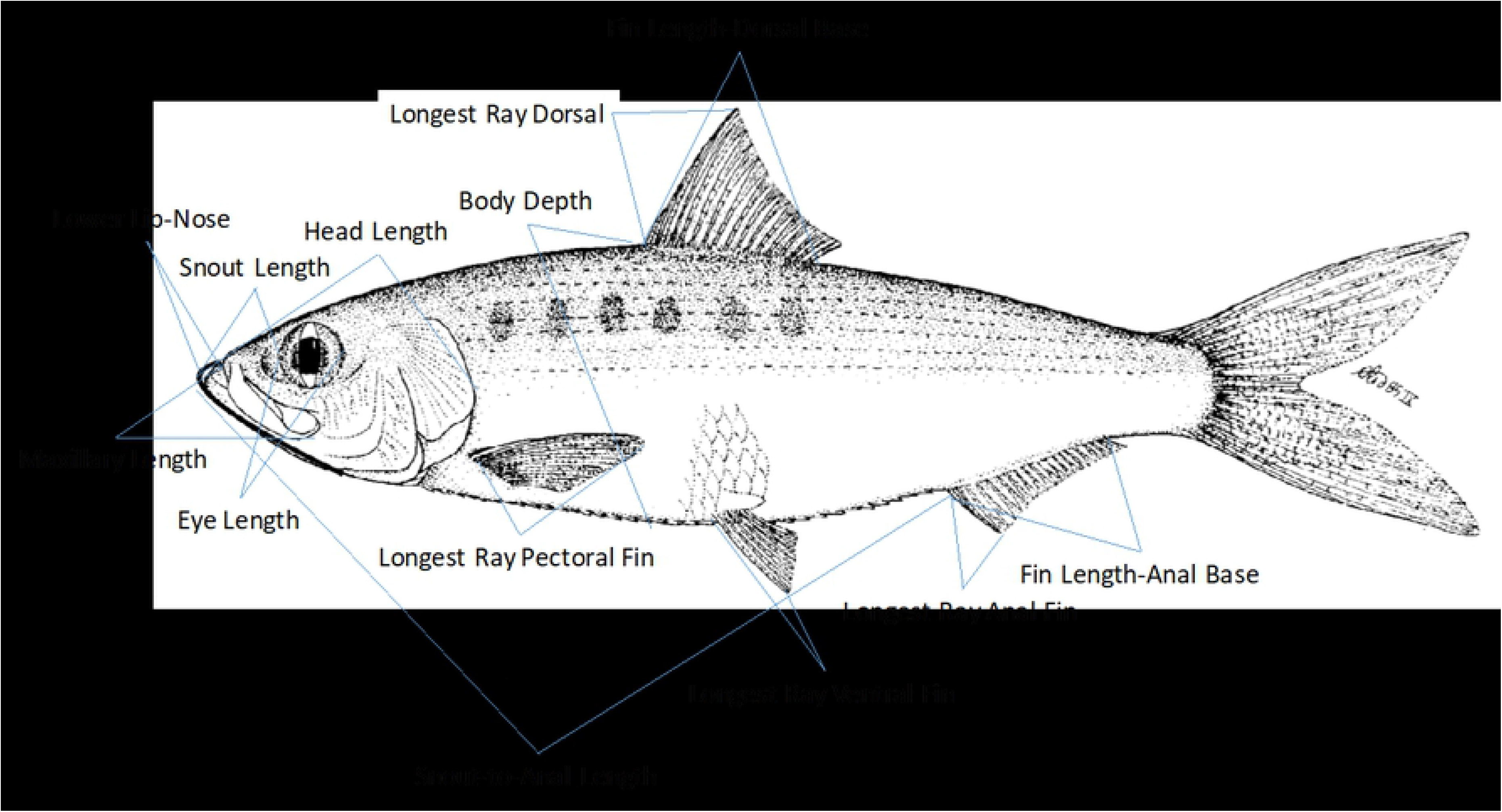
Hickory Shad illustration showing how morphometric measurements were taken. Reproduced from Whitehead [34].

External meristic counts were taken on the left side of the body, unless there was damage: post ventral (pelvic) scutes (PVS) -- count of scutes from the end of the ventral fin to the anus; anterior ventral scutes (AVS) -- count of scutes from the beginning of the operculum to the ventral fin, including the scute straddling the ventral fins; scale rows (SR) -- count of scales along the lateral line, beginning at the upper angle of the operculum and terminating at the end of the hypural plate as determined with a crease in the caudal peduncle by folding the tail; and longitudinal scale rows (LSR) -- count of scales from the origin of the dorsal fin to the origin of the ventral fin. A random subset of specimens (n = 463) were analyzed for an additional four internal meristic counts, including the left and right gill rakers of the upper first arch (L-GRU, R-GRU) -- count of all gill rakers on the upper arch of first gill raker, not including the raker straddling the angle; and left and right gill rakers lower (L-GRL, R-GRL) -- count of all first arch gill rakers from the raker straddling the angle to the end, regardless of size.

External meristic characters including the scale rows between the upper angle of gill opening and base of caudal fin, longitudinal scale rows between origin of ventral fin and origin of dorsal fin, post-ventral scutes, and anterior-ventral scutes, were all counted from the freshly-thawed specimens.

To the best of our knowledge, there are no references in the literature detailing specific methods for counting scutes of clupeids. We chose to divide the scute count into two -- anterior and posterior -- of the ventral fin following Smith [17], though Nichols [26] and Melvin et al. [19] chose to count total scutes for American Shad. All scutes were counted, regardless of size, from where the ventral surface reaches the operculum posterior to the anus. Special care was given to check for scutes obscured by the anus in all fish, specifically ripe females. Occasionally scales near the scutes had to be removed to fully expose all scutes, and then counts were obtained with the aid of a probe.

After external morphometric measurements and meristic counts were completed, fish were then dissected to remove the gonads, which were weighed to the nearest 0.01 g. Sex was determined for each specimen based on visual inspection of the gonad. Once features of each specimen were recorded, the data were compiled into one Microsoft Excel file for analyses.

Sample sizes for each state, watershed, and capture location were not uniform, nor were the number of males and females the same, due to the various collection methods and availability at the time of collection. In addition, the number of fish analyzed for each character was not always equal because some of the specimens were damaged necessitating the omission of one or more characters. Also the timing of the collection for each watershed was not standardized; spawning often started prior to the typical timeline for state agency spring sampling. The morphometric and meristic data presented here are from frozen and thawed -- not fresh -- Hickory Shad and for purposes of the analyses we assumed that any bias caused by this process was equal across all specimens.

## Results

Overall 717 Hickory Shad were analyzed for 17 morphometric measurements and four meristic characters from 23 different rivers and estuaries in Maryland, Delaware, Virginia, North Carolina, South Carolina, Georgia, and Florida following the methods outlined above. Results of descriptive statistics for all locations combined, separated by sex, for all measurements and counts are presented in Table 3. Results for each individual river and combined sex can be found in Table 4. The random subset of specimens (n = 463) analyzed for four internal meristic gill raker counts showed that Hickory Shad had between 8-11 rakers on L-GRU, 8-12 rakers on R-GRU, and 19-22 rakers on both L-GRL and R-GRL.

**Table 3.** Descriptive data of morphometric and meristic characters for female and male specimens of Hickory Shad. See text for descriptions of each measurement or count. All measurements given in mm.

| Female |  |  |  |  |  | Male |  |  |  |  |  |
| --- | --- | --- | --- | --- | --- | --- | --- | --- | --- | --- | --- |
| Character | Range | Mean | SD | % SL | n | Character | Range | Mean | SD | % SL | n |
| SL | 229 - 389 | 292.41 | 26.09 | - | 365 | SL | 206 - 344 | 264.62 | 20.63 | - | 330 |
| FL | 260 - 435 | 326.36 | 28.57 | 111.61 | 365 | FL | 232 - 382 | 295.81 | 22.64 | 111.78 | 330 |
| TL | 306 - 508 | 383.36 | 33.55 | 131.10 | 364 | TL | 272 - 444 | 347.67 | 26.48 | 131.38 | 326 |
| LLN | 2.44 - 7.90 | 3.93 | 0.73 | 1.34 | 365 | LLN | 2.39 - 7.39 | 3.60 | 0.54 | 1.36 | 327 |
| SAL | 172 - 289 | 218.99 | 20.26 | 74.89 | 364 | SAL | 155 - 251 | 197.51 | 15.45 | 74.64 | 330 |
| BD | 65.74 - 134.89 | 91.44 | 13.32 | 31.27 | 365 | BD | 60.70 - 105.09 | 79.03 | 7.46 | 29.86 | 329 |
| HL | 64.73 - 108.43 | 81.40 | 6.96 | 27.84 | 364 | HL | 58.86 - 93.96 | 74.72 | 5.99 | 28.24 | 329 |
| EL | 11.82 - 19.60 | 14.80 | 1.26 | 5.06 | 363 | EL | 11.53 - 18.59 | 13.96 | 1.16 | 5.28 | 330 |
| SNL | 15.93 - 27.84 | 20.75 | 1.84 | 7.10 | 363 | SNL | 15.13 - 24.76 | 19.07 | 1.63 | 7.21 | 330 |
| HW | 23.67 - 47.07 | 31.33 | 3.38 | 10.71 | 363 | HW | 21.87 - 37.11 | 28.45 | 2.53 | 10.75 | 330 |
| IOW | 10.38 - 20.90 | 14.52 | 1.77 | 4.96 | 364 | IOW | 9.56 - 20.75 | 13.35 | 1.60 | 5.05 | 329 |
| ML | 26.51 - 41.90 | 33.98 | 2.52 | 11.62 | 362 | ML | 25.38 - 37.15 | 31.51 | 2.18 | 11.91 | 330 |
| FLD | 33.97 - 63.41 | 63.41 | 4.80 | 21.69 | 365 | FLD | 29.66 - 52.89 | 39.77 | 3.74 | 15.03 | 329 |
| FLA | 37.80 - 68.23 | 49.00 | 4.65 | 16.76 | 363 | FLA | 32.43 - 62.17 | 44.65 | 4.15 | 16.87 | 328 |
| LRD | 29.88 - 55.12 | 39.77 | 3.99 | 13.60 | 364 | LRD | 24.63 - 45.91 | 36.23 | 3.32 | 13.69 | 327 |
| LRP | 43.10 - 77.09 | 55.42 | 5.32 | 18.95 | 363 | LRP | 38.84 - 64.67 | 50.83 | 4.45 | 19.21 | 330 |
| LRV | 23.76 - 46.09 | 34.90 | 3.29 | 11.93 | 362 | LRV | 23.80 - 39.84 | 31.97 | 2.88 | 12.08 | 330 |
| LRA | 14.61 - 26.59 | 19.74 | 2.33 | 6.75 | 360 | LRA | 13.13 - 23.60 | 17.99 | 1.89 | 6.80 | 326 |
| SR | 49 - 56 | 51.65 | 1.23 |  | 349 | SR | 49 - 56 | 51.60 | 1.12 |  | 316 |
| LSR | 15 - 19 | 17.81 | 0.55 |  | 339 | LSR | 15 - 19 | 17.71 | 0.60 |  | 309 |
| PVS | 14 - 18 | 15.86 | 0.75 |  | 361 | PVS | 14 - 18 | 15.88 | 0.72 |  | 326 |
| AVS | 17 - 23 | 21.35 | 0.84 |  | 363 | AVS | 18 - 24 | 21.39 | 0.79 |  | 329 |

**Table 4.**
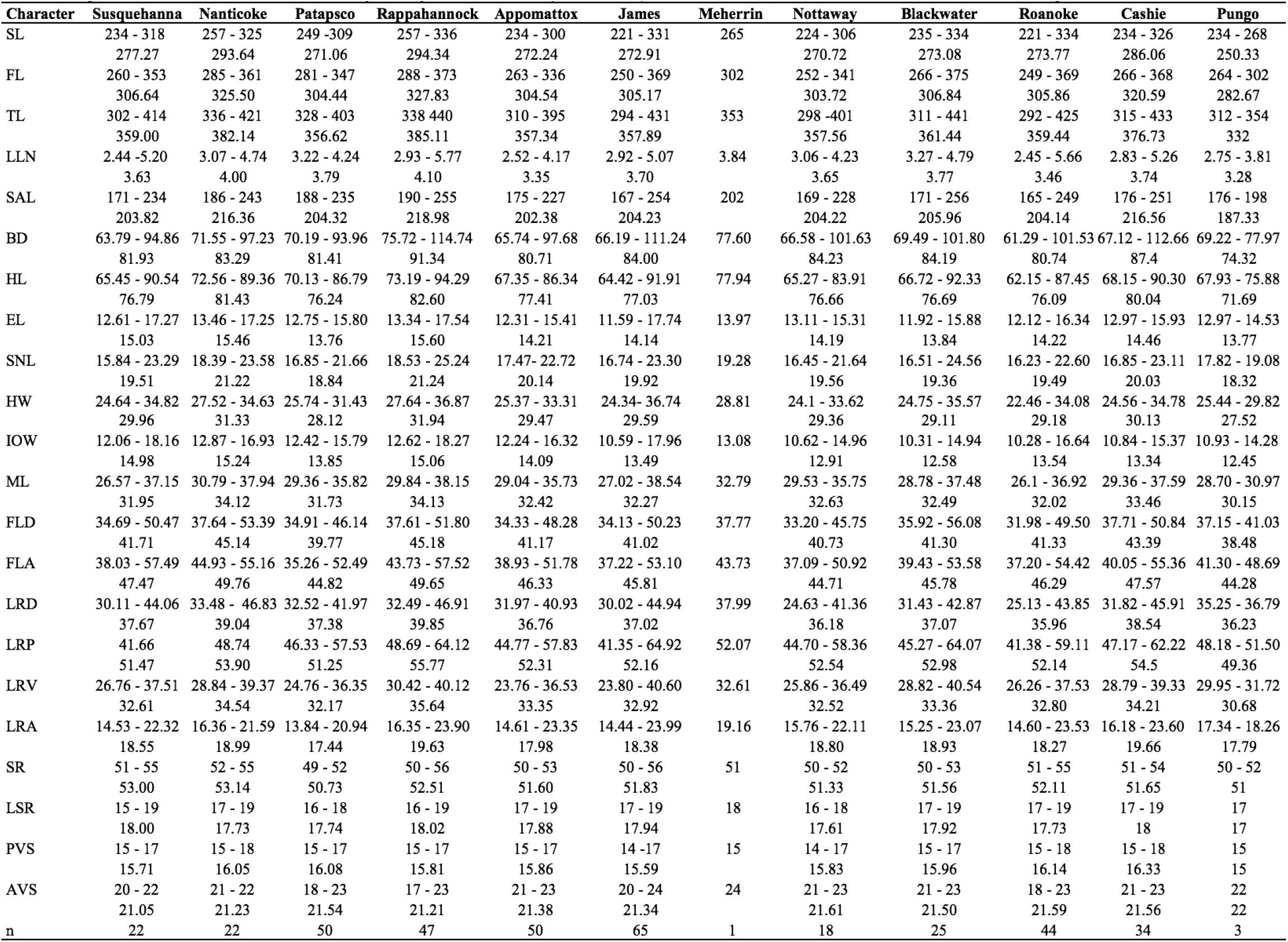
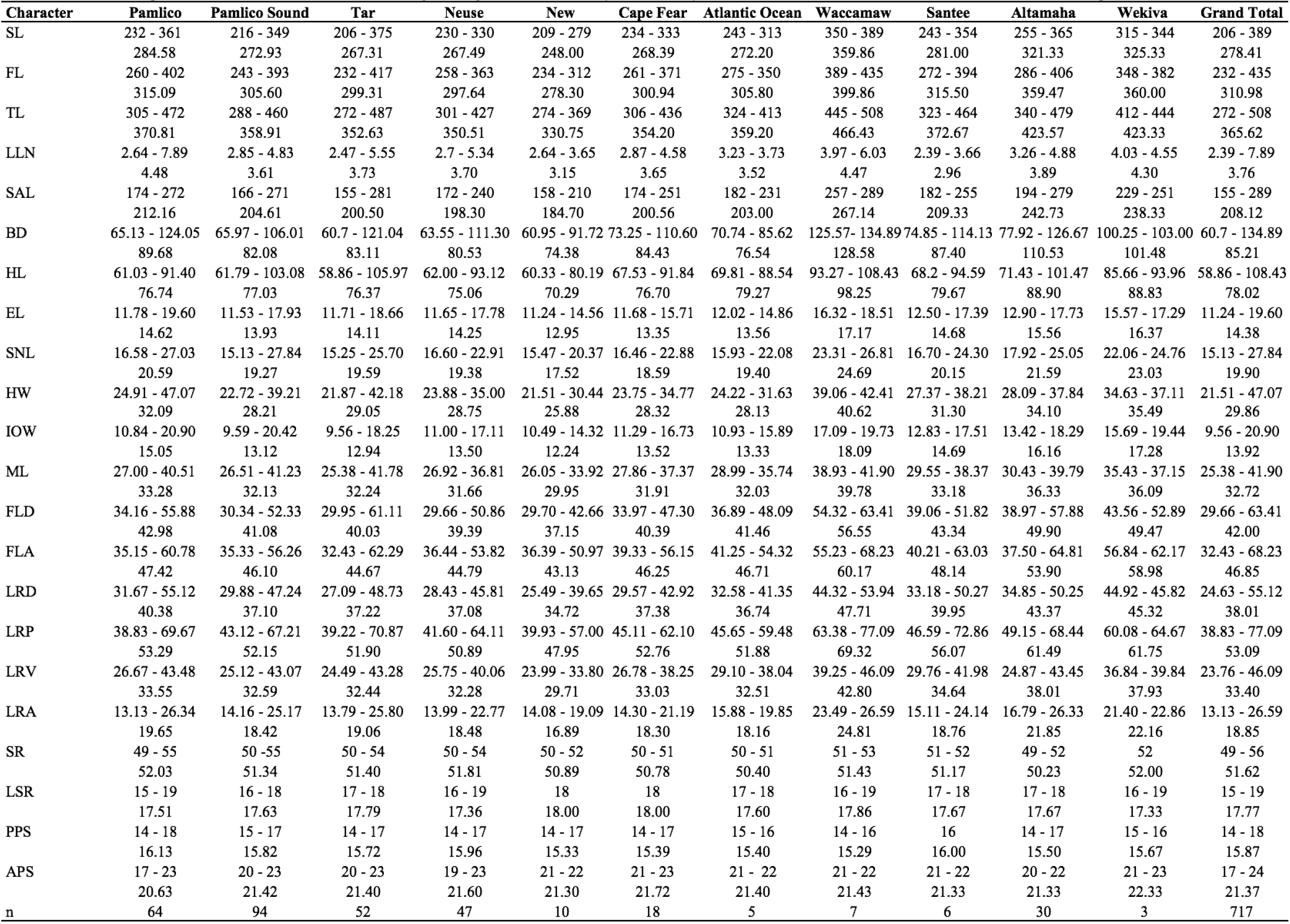
Morphometric and meristic charcters of Hickory Shad by river of collection (north to south). For each character; minimum - maximum, and mean values are presented.

A basic review of the morphometric and meristic data showed sexual difference in many characters, namely morphometric measurements. All morphometric characters showed sexual difference in character means, yet some character differences were more substantial. For example, the mean measurements (mm) of BD (Female: 91.44, Male: 79.03), FLD (Female: 63.41, Male: 39.77), SAL (Female: 218.99, Male: 197.51), and HL (Female: 81.40, Male: 74.72) were largely different between sexes. For meristic counts on SR, LSR, PVS, and AVS there was no observed difference between sexes and so the averages between males and females were similar. Of the four counts, the largest difference in the averages was found for the count of LSR where the averages were 17.81 and 17.71 for females and males, respectively. Due to the differences in some characters (i.e., morphometric) by sex, it was necessary to divide the morphometric and meristic data for males and females for accurate description and analysis.

### Specimen Size

All Hickory Shad collected and included in this study were adults (sexually mature) participating in the annual spawning run and all morphometric and meristic data reported are for adult fish. Male specimens from locations combined ranged from 206 to 344 mm SL with a mean + SD of 264.42 + 20.52 mm. Female specimens ranged from 229 to 389 mm with a mean of 289.72 + 24.71 mm. Sizes for sexes and locations combined ranged from 206 to 389 mm SL with a mean of 276.53 + 26.31 mm. The linear relationships between FL and TL, and FL and SL, were:

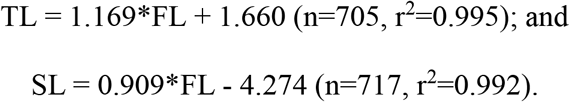

The largest Hickory Shad were from the Waccamaw River, SC and the mean + SD was 359.86 + 13.92 mm SL with a range between 350 and 389 mm SL; average weight was 1281.13 + 95.46 g and all seven specimens from this river were female. On average the smallest Hickory Shad were collected from the New River, NC with a mean + SD of 248 + 24.92 mm SL and a range of 209 to 279 mm SL. However, the smallest Hickory Shad collected in this study (206 mm SL) was a male from the Tar River, NC. Specimen total body weight (n = 695) with sexes and locations combined ranged from 206.03 to 1488.28 g with a mean + SD of 501.34 + 187.52 g, and gonad weights (n = 691) from 0.38 to 266.03 g with a mean of 49.88 + 45.78 g.

### Sex

Overall, females were larger than males of similar SL. The smallest male weighed 206.03 g and largest weighed 866.50 g. The smallest female weighed 242.40 g and largest 1488.28 g. Gonads for females weighed from 0.38 to 266.03 g with a mean + SD of 74.43 + 51.15 g. Gonads for males weighed from 0.90 to 62.53 g with a mean of 22.86 + 11.53 g. Variation in size and weight of female gonads were largely dependent on spawning status. Some gonad specimens had deteriorated so gonad weight measurements (n = 4) and sex determination (n = 22) were not possible. In addition, the sexing of some specimens was omitted on the data sheet during the examination process.

### Missing data

Some of the 717 Hickory Shad could not be analyzed for the entire suite of 17 morphometric and four meristic characters due to specimen damage. This resulted in 146 missing values across all morphometric and meristic characters. Missing value analysis was performed in SPSS version 24 [27] and the meristic character LSR had the most missing data (7.7%). Of the remaining characters only SR, LRA, and PVS had more than 1.0% missing: 4.3, 1.3, and 1.1%, respectively. Values for count and percent missing of each character are reported in Table 5.

**Table 5.**
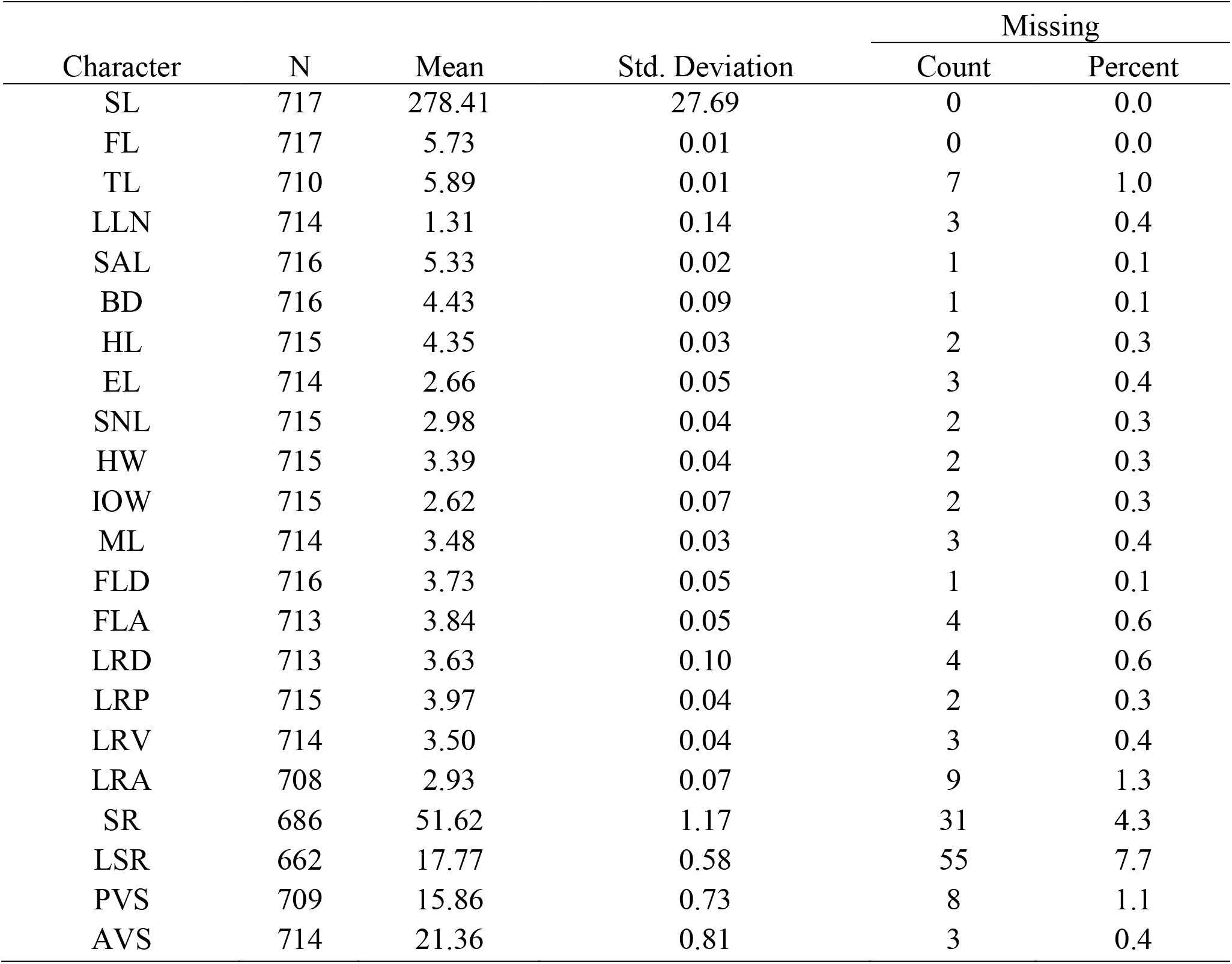
Missing value analysis of 18 morphometric and four meristic characters of Hickory Shad.

| Character | N | Mean | Std. Deviation | Missing |  |
| --- | --- | --- | --- | --- | --- |
|  |  |  |  | Count | Percent |
| SL | 717 | 278.41 | 27.69 | 0 | 0.0 |
| FL | 717 | 5.73 | 0.01 | 0 | 0.0 |
| TL | 710 | 5.89 | 0.01 | 7 | 1.0 |
| LLN | 714 | 1.31 | 0.14 | 3 | 0.4 |
| SAL | 716 | 5.33 | 0.02 | 1 | 0.1 |
| BD | 716 | 4.43 | 0.09 | 1 | 0.1 |
| HL | 715 | 4.35 | 0.03 | 2 | 0.3 |
| EL | 714 | 2.66 | 0.05 | 3 | 0.4 |
| SNL | 715 | 2.98 | 0.04 | 2 | 0.3 |
| HW | 715 | 3.39 | 0.04 | 2 | 0.3 |
| IOW | 715 | 2.62 | 0.07 | 2 | 0.3 |
| ML | 714 | 3.48 | 0.03 | 3 | 0.4 |
| FLD | 716 | 3.73 | 0.05 | 1 | 0.1 |
| FLA | 713 | 3.84 | 0.05 | 4 | 0.6 |
| LRD | 713 | 3.63 | 0.10 | 4 | 0.6 |
| LRP | 715 | 3.97 | 0.04 | 2 | 0.3 |
| LRV | 714 | 3.50 | 0.04 | 3 | 0.4 |
| LRA | 708 | 2.93 | 0.07 | 9 | 1.3 |
| SR | 686 | 51.62 | 1.17 | 31 | 4.3 |
| LSR | 662 | 17.77 | 0.58 | 55 | 7.7 |
| PVS | 709 | 15.86 | 0.73 | 8 | 1.1 |
| AVS | 714 | 21.36 | 0.81 | 3 | 0.4 |

### Comparison between Hickory Shad and other Clupeids

Morphometric and meristic results of this study were compared to available literature values for morphologically similar clupeids, including the American Shad, Alewife, and Blueback Herring (Table 6). Characters mentioned here represent the clearest difference between species: Hickory Shad have a larger body depth as a percent of total length (22.31-26.55) compared to American Shad (17.2-19.4) and Alewife (17.8-21.7), but body depth is similar to that of Blueback Herring (22.1-25.2). The upper portions of the variable ranges for Hickory Shad scute and scale row counts (PVS, AVS, and SR) were less than that for American Shad, but LSR was greater for Hickory Shad. This is not surprising since the body depth as a percent of total length was greatest for Hickory Shad, and the LSR character is counted along the depth of the body. The range of interorbital width (IOW) as a percent of head length for Hickory Shad (16.24-19.28) was most similar to Alewife (15.7-21.6); the range for American Shad (18.6-21.6) was higher than for Hickory Shad but within the range for Alewife. Overall, Blueback Herring interorbital width as a percent of head length (21.1-26.4) is the largest. As for eye length as a percent of head length, the Hickory Shad has the smallest range (18.08-19.10), which is much less than that of American Shad (27.3-32.0), Blueback Herring (23.4-30.0) and Alewife (26.9-35.7).

**Table 6.** Comparison of morphometric and meristic characters for Hickory Shad, American Shad, Alewife, and Blueback Herring. Range given for each, if available; usual values reported in literature in parentheses.

| Character | Hickory Shad | American Shad | Blueback Herring | Alewife |
| --- | --- | --- | --- | --- |
| BD % TL | 22.31-26.55 | 17.2-19.4 <sup>a</sup> | 22.1-25.2 <sup>a</sup> | 17.8-21.7 <sup>a</sup> |
| HL % TL | 21.34-21.64 | 22.7-24.0 <sup>a</sup> | 18.5-20.6 <sup>a</sup> | 20.3-23.7 <sup>a</sup> |
| EL % HL | 18.08-19.10 | 27.3-32.0 <sup>a</sup> | 22.0-26.4 <sup>a</sup> | 26.1-32.0 <sup>a</sup> |
| SNL % HL | 25.68-25.71 | 26.9-32.0 <sup>a</sup> | 23.4-30.0 <sup>a</sup> | 26.9-35.7 <sup>a</sup> |
| IOW % HL | 16.24-19.28 | 18.6-21.6 <sup>a</sup> | 21.1-26.4 <sup>a</sup> | 15.7-21.6 <sup>a</sup> |
| FLA % TL | 11.92-13.43 |  |  | 10.3-12.0 <sup>a</sup> |
| PVS | 14-18 | 12<sup>b</sup>-19<sup>a</sup> | 12-16 <sup>a</sup> | 12<sup>d</sup>-17<sup>f</sup> (14-15)<sup>a</sup> |
| AVS | 17-24 | 19-25 <sup>b</sup> | 18-21 <sup>d</sup> | 17-21 (19-20) <sup>a</sup> |
| SR | 49-56 | 52-64 <sup>c</sup> | 46-54 <sup>d</sup> | 42<sup>d</sup>-54<sup>g</sup> |
| LSR | 15-19 | 15-16 <sup>d</sup> | 13-14 <sup>e</sup> | 14 <sup>d</sup> |

<sup>a</sup> Scott and Crossman 1973, <sup>b</sup> Hill 1956, <sup>c</sup> Walburg and Nichols 1967, <sup>d</sup> Hildebrand 1963
<sup>e</sup> Thomson et al 197, <sup>f</sup> Leim and Scott 1966, <sup>g</sup> Miller 1957

## Discussion

It is often difficult to discern the causes of morphological and meristic variations between fish populations [22] though it is assumed they might be related to genetic differences or linked to phenotypic plasticity resulting from non-homogeneous environmental factors in each river [19]. However, reasons why there are variations in meristic and morphological characters were not an objective of our study.

Instead, our study provides foundational information on the morphometric and meristic variation of Hickory Shad across a large portion of the species range. To complement this study, further research is needed to investigate these characters of Hickory Shad from more southern rivers in Georgia and Florida. This would allow comparison of morphometric and meristic variation across the entire species range and determine if greater geographic distance corresponds to larger variation. It is more likely that adjacent rivers or watersheds share common environmental characteristics compared to rivers separated by large distances, possibly leading to greater variation in morphometrics and meristics. For instance, we were able to obtain 22 samples from a small tributary of the Susquehanna, River Maryland at the mouth of Deer Creek (39.613358 N, −76.149024 W), which is near the northern end of the assumed Hickory Shad spawning range. Unfortunately, we were unable to obtain large sample sizes from the southernmost Hickory Shad spawning population of the St. Johns River, Florida, though we did obtain three specimens from the Wekiva River, a tributary of the St. Johns (28.8728226 N, - 81.3689402 W). The Wekiva River, Florida, and Deer Creek, Maryland, are separated by roughly 1280 Km.

One limitation of this study is that equal sample sizes for each state and watershed could not be collected. Attempts were made to have between 25-50 fish per watershed and a 50:50 sex ratio, but as with most all fisheries work, success in sampling is often not reliable. Multiple factors influenced our ability to collect more samples, including early Hickory Shad spawning runs in some locations, foul weather, low river water levels prohibiting boat access, severe long-term flooding, and expense of traveling to distant locations. It is possible that the morphometric and meristic values presented here for rivers with small samples sizes may not accurately capture the true natural variation of the characters in those populations. Additionally, the timing of specimen collection was not standardized and often started after the spawning run had fully begun, which could have potentially affected this study (i.e., size or sex distributions). Overall, slightly more female specimens (n = 365) were collected than male (n = 330) representing 52.5% and 47.5% of the specimens included in this study, respectively. The difference in the number of males and females could be a product of gear bias and not necessarily representative of the natural populations. For instance, gill nets used to collect some specimens in this study are more selective for larger female Hickory Shad than smaller males. Melvin et al. [19] studying American Shad also found gill nets to be selective for larger females. Furthermore, we experienced a willingness of sport fishers to provide specimens for our study, but reluctance to provide females since most fishers wanted the roe for bait or for personal consumption.

### Sexual Differences

Differences observed in the averages of morphometric characters when compared by sex was not a surprising result and is relatively common in fish, though it has never been explicitly described for Hickory Shad. This has significant implications and suggests studies on Hickory Shad must be separated by sex and analyzed in that manner since there is substantial difference between male and female specimens. Melvin et al. [19] came to similar conclusions for morphometric and meristic characters of American Shad and so males and females were analyzed separately.

### Specimen Size

It is important to note that the morphometric measurements presented in this study are of frozen and not freshly caught Hickory Shad. It is possible that the freezing and thawing process may slightly alter the shape and or size of some morphometric characters. Melvin et al. [28] reported a significant difference (P < 0.01) between length measurements of live American Shad in the field compared to measurements of dead specimens in the laboratory. In the event American Shad were frozen prior to measurement, the length was multiplied by 1.021 to better approximate fresh length [28]. Though fish samples are often frozen by biologists for later processing, future studies should investigate if there is a significant difference between morphometric measurements for fresh versus frozen Hickory Shad and, if so, which measurements are the most robust to the freezing and thawing process. Cronin-Fine et al. [29] found 10 geometric morphometric measurements of Alewife that did not have a significant difference between fresh and frozen specimens. Generally for meristics, the act of freezing and thawing is not problematic since it does not change the counts of meristic features.

The freezing and thawing process could also have biased the weight of the fish, but similar to morphometric measurements, the bias is shared across all individuals. Also, gonad weight can be extremely dependent on spawning status (pre or post-spawn), especially for females. Spent females weigh less than ripe and ready-to-spawn individuals, but unfortunately spawning status was not recorded during dissections. There were a few instances of gonads that were unable to be weighed (or sexed) because they were no longer intact or starting to decompose. This was likely a result of freezer storage for an extended length of time, multiple freezing and thawing events, or the length of time from collection till initial freezing. This was not a serious problem; 26 specimens exhibited deterioration and this state was relatively random across rivers. Also, it was likely that some of the individuals not sexed was caused by human error instead of relating to the state of the gonads.

The regression equations for relationships between Hickory Shad FL and TL, and between FL and SL, provide a means for converting between the various measurements of fish size. This could be useful for biologists or fishery managers to accurately estimate one length from another in the instance that only one of the measurements was recorded.

### Missing Data

Though not a frequent problem in this study, missing data are quite common in morphometric (and meristic) studies [30]. Some of the specimens could not be analyzed for the entire 17 morphometric and four meristic characters due to damage including broken or missing fins, missing scales, and wounds from predation or gear-related injury. Missing scales are not surprising, since the Hickory Shad as well as other clupeids are very susceptible to shedding scales. The frequency of missing values for all characters can be found in Table 5. In our study no imputation procedures (i.e., replacement or regression-based approaches) were used to estimate missing data; instead these values were simply omitted.

### Comparison between Hickory Shad and other Clupeids

Most of the morphometric and meristic characters investigated in this study do not serve to easily differentiate Hickory Shad from American Shad, Alewife, or Blueback Herring though careful examination of certain characters can help narrow down the species. One common and definitive way to distinguish Hickory Shad from the other species is by gill raker counts. Though not directly incorporated into this study, a random subset of Hickory Shad specimens was analyzed for gill raker counts. It was determined that Hickory Shad had between 19-22 gill rakers on the lower limb of the first arch (n=463), which is considerably less than the other anadromous *Alosa* species. American Shad typically have 59-76 lower gill rakers on the first arch, Blueback Herring 41-52, and Alewife 38-46, all of which are higher counts [31] due to their diet being different than Hickory Shad, which are more piscivorous [32].

## Conclusion

Mansueti [7] described Hickory Shad as “The most enigmatic of all estuarine clupeoids” and the intent of our study was to expand the existing taxonomic knowledge of the species. Mitchill [2] used six meristic characters in describing the species: branchiostegal, pectoral, ventral, anal, dorsal, and caudal rays. These six characters were not included in this study because the methods Mitchill used to count them were not available and therefore no direct comparison was possible. Instead, 17 morphometric measurements and four meristic counts not included in the original description of the species were utilized. The information about the anatomical characteristics presented herein are lacking in the literature, though they are well known for most other anadromous fish species. These additional morphological and meristic characters may prove valuable for separating regions or watersheds in future studies. Geometric morphometric analysis may be another viable option to investigate body shape variability. In addition, there still remain many unanswered questions regarding Hickory Shad life history, biology, and stock status that should be addressed so that the species can be properly managed and all spawning populations sustained. Furthermore, the intraspecific variation of Hickory Shad described here could be used to discriminate the different populations using multivariate analysis [33].

## Acknowledgments

We thank the North Carolina Wildlife Resources Commission for providing funding through the Sport Fish Restoration Act. We also thank the many agencies that collected samples: North Carolina Wildlife Resources Commission, North Carolina Division of Marine Fisheries, South Carolina Department of Natural Resources, Florida Fish and Wildlife Conservation Commission, Virginia Department of Game and Inland Fisheries, Maryland Department of Natural Resources, Georgia Department of Natural Resources, Delaware Division of Fish and Wildlife, and the Smithsonian Environmental Research Center. The Department of Biology at East Carolina University provided logistic support, travel, and field and laboratory equipment.

## Author Contributions

**Conceptualization**: Roger A. Rulifso

**Data curation:**: Roger A. Rulifson

**Formal analysis**: Jordan P. Smith; Michael W. Brewer

**Funding acquisition**: Roger A. Rulifson.

**Investigation**: Jordan P. Smith

**Methodology**: Jordan P. Smith, Michael W. Brewer, Roger A. Rulifson

**Project administration**: Roger A. Rulifson.

**Resources**: Roger A. Rulifson.

**Software**: Jordan P. Smith

**Supervision**: Roger A. Rulifson, Michael W. Brewer

**Visualization**: Jordan P. Smith

**Writing - original draft**: Jordan P. Smith

**Writing - review & editing**: Roger A. Rulifson, Michael W. Brewer, Jordan P. Smith

